# IS110 transposon utilizes two mechanistically distinct RNA-guided transposition pathways

**DOI:** 10.1101/2025.10.25.684509

**Authors:** Xuanlong Sun, Chen Yang, Zhaohui Cai, Li Liu, Xinmin Yue, Shuquan Rao, Huiqiang Lou, Chaoyou Xue

## Abstract

The recently discovered IS110 elements employ RNA-guided transposases that use bridge-RNA to coordinate donor and target DNA recognition and integration via donor- and target-binding loops, offering promise for precise, DSB-free genome editing. However, their transposition mechanism remains unclear, with prior inconsistent observations. Using a IS110 element from *Caloranaerobacter azorensis*, we show that IS110 transposons utilize two independent transposition pathways. The first depends on bridge-RNA and excises the element via copy-out to form a circular intermediate, revisiting the earlier cut-out model. Integration occurs predominantly through top-strand insertion, suggesting an alternative to the earlier Holliday junction-mediated dual-strand mechanism. The second involves cooperation between a truncated target-binding loop RNA and bridge-RNA, enabling direct top-strand transfer to a new target via two sequential reactions, bypassing circular intermediates. Both pathways preserve the original transposon copy. These findings redefine the mechanistic understanding of IS110 transposition and reconcile prior discrepancies, offering insight into programmable, RNA-guided genome modification.

## Introduction

Insertion sequence (IS) elements, the simplest transposable elements, encode only a transposase that mediates excision and integration into new genomic loci, enabling their propagation^1^. Transposases recognize IS ends through sequence-specific protein-DNA interactions to facilitate transposition^2^. Unlike CRISPR-Cas systems, which use guide RNAs for programmable targeting^3,4^, transposase-based systems rely on fixed protein-DNA interactions, limiting their programmability.

Recent studies by Durrant et al. and Siddiquee et al. uncovered a novel RNA-guided transposon system, where the IS110 transposase employs a compact bridge-RNA (bRNA) to coordinate donor and target DNA recognition and integration^5-7^. The bRNA contains two loops, a target-binding loop (TBL) with left and right target guide regions (LTG and RTG) that base-pair with the bottom strand of the left target (LT) and the top strand of the right target (RT), and a donor-binding loop (DBL) with left and right donor guide regions (LDG and RDG) that hybridize to the bottom strand of the left donor (LD) and the top strand of the right donor (RD). This system was hypothesized to follow a cut-out-paste-in mechanism^6,8,9^, where the transposon is excised as a circular dsDNA intermediate by rejoining LT-RD and LD-RT, restoring the donor (LD-RD), followed by integration at new target sites. However, non-regenerated target sites observed by Siddiquee et al^7^. suggest an alternative mechanism. For integration, while Durrant et al. and Hiraizumi et al. proposed a bRNA-dependent Holliday junction mechanism^5,6^, Siddiquee et al. showed that TBL-only RNA can mediate transposition^7^, raising critical questions: how is donor DNA recognized without DBL? Does the TBL indirectly recruit donor DNA through uncharacterized interactions? These discrepancies highlight critical mechanistic gaps.

Despite these gaps (Fig. S1A), the IS110-bRNA system offers transformative potential for double-strand break (DSB)-free genome editing. Unlike CRISPR-Cas systems, which rely on cellular DNA repair and perform poorly in non-dividing cells^3,10^, IS110-bRNA system enables precise integration without DSBs. It also shows strong potential for inserting large DNA fragments, a challenge for CRISPR-Cas systems due to inefficient repair of large inserts^11^. To harness this potential, understanding IS110 transposition mechanism is crucial. Here, using a IS110 element from *Caloranaerobacter azorensis*, we show that IS110 transposons employ two independent transposition pathways. The first resembles a bRNA-depended copy-out-paste-in mechanism, involving top-strand excision and rejoining, followed by resolution into circular ssDNA or dsDNA intermediates. Both forms can be integrated, but dsDNA integrates predominantly via top-strand, with >1,300-fold higher efficiency than bottom-strand, implying additional factors are involved in resolving the initial single-strand insertion into a double-stranded product. The second pathway bypasses circular intermediates, where the transposase, sequentially transfers LT-RD and LD-RT directly to target sites. While TBL-only RNA supports partial transposition via LT-RD transfer, complete cargo mobilization requires full-length bRNA for the LD-RT step. This RNA-guided direct transfer mechanism represents a non-canonical form of transposition.

## Results

### *Caz*IS110-1 is active for excision and integration

To select a suitable IS110 transposase for mechanistic studies, we screened for transposons from thermophilic organisms (for improved protein stability) with high genomic copy numbers (indicating activity). We selected *Caz*IS110-1 from the thermophilic bacterium *Caloranaerobacter azorensis* (*Caz*)^12^, which has 10 genomic copies (Fig. 1A). *Caz*IS110-1 transposase was purified with its predicted bRNA. In the absence of bRNA, *Caz*IS110-1 precipitated, but MBP fusion rendered it soluble and catalytically active, enabling *in vitro* assays (Fig. S1B). This bRNA forms two loops: TBL with LTG and RTG, and DBL with LDG and RDG, which base-pair with the LT-RD and LD-RT junctions of the *Caz*IS110-1 transposon, respectively (Fig. 1B).

**Figure 1.**
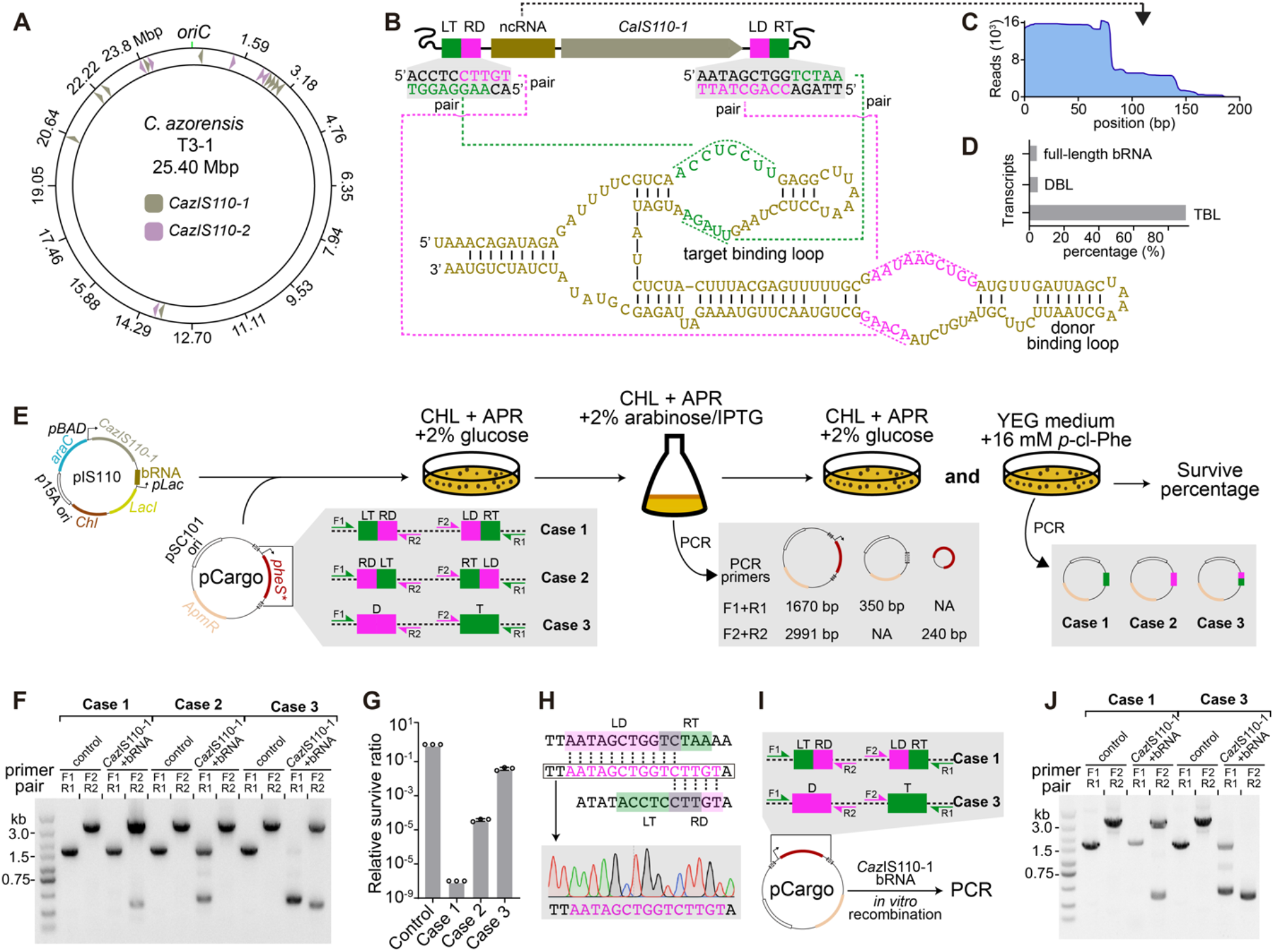
RNA-guided transposition of *Caz*IS110-1. **A**, Genome-wide distribution of *Caz*IS110-1 and *Caz*IS110-2 elements in *Caloranaerobacter azorensis* strain T3-1. **B**, Predicted bRNA secondary structure of *Caz*IS110-1 element. Sequences pairing with LT-RD and LD-RT junctions of the *Caz*IS110-1 transposon are highlighted in green and magenta, respectively. **C**, Alignment of small RNA-seq reads to the predicted bRNA sequence. **D**, Relative abundance of TBL-only, DBL-only and full-length bRNA species. **E**, Schematic of three *in vivo* transposition assays: Case 1, *phes*\* is inserted between LT-RD and LD-RT junctions; Case 2, *phes*\* is inserted between LD-RT and LT-RD junctions; Case 3, *phes*\* is inserted between donor and target sites. Successful transposition enables growth on YEG medium supplemented with *p*-Cl-Phe. Primer pairs and expected amplicons are indicated. **F**, PCR detection of *in vivo* transposition products from **E. G**, Quantification of *in vivo* transposition efficiency from **E. H**, Sanger sequencing of the PCR products confirming donor site rejoining. **I**, Schematic of *in vitro* transposition assay. **J**, PCR detection of *in vitro* transposition products. Data in **G** are mean ± s.d. for n=3 biologically independent experiments.

To precisely identify RNAs associated with *Caz*IS110-1, we extracted RNA from affinity-purified RNP complexes and performed RNA sequencing. Alignment of sequencing reads to the predicted bRNA revealed multiple transcripts (Fig. 1B-D). Among these, we identified a ∼170 nt transcript corresponding to the predicted full-length bRNA, a ∼77 nt transcript containing only the TBL, and an 87 nt transcript containing only the DBL (Fig. 1C and S1C). The ∼77 nt TBL-only RNA transcript is the predominant RNA species, comprising 90.4% of transcripts. The striking abundance of TBL-only RNA suggests potential functional divergence from the bRNA (Fig. 1D).

We next investigated the role of bRNA, TBL-only and DBL-only RNAs in transposition, we developed a plasmid-based transposition assay in *E. coli*, using a *phes*\* counterselection system^13^. Excision or integration removes *phes*\*, enabling *E. coli* survival on YEG medium supplemented with *p*-cl-Phe (Fig. 1E). We constructed an expression plasmid encoding *Caz*IS110-1 and the RNAs (pIS110-1), alongside target plasmids containing either *phes*\* flanked LT-RD/LD-RT junctions (for excision) or target/donor sites (for integration) (Fig. S1D). Following induction, we performed PCR on cellular lysates to detect excised transposon circles and target-donor integration joints. In excision assays, we observed PCR products corresponding to the rejoined donor site (LD-RD) (Fig. 1F). In integration assays, PCR products representing target-donor integration joints (LT-RD and LD-RT) were detected (Fig. 1F). No PCR products were observed without *Caz*IS110-1 or bRNA, or when truncated RNAs (TBL- or DBL-only) were used, confirming that both excision and integration require the full-length bRNA (Fig. S1E). Consistent with previous studies, target-site regeneration was absent (Fig. 1E). Colony PCR and Sanger sequencing of cells on selective media quantified excision (∼10^-4^) and integration (∼10^-1^) efficiencies (Fig. 1G), with sequencing confirming donor-site rejoining and LT-RD/LD-RT integration (Fig. 1H and S1F). Colonies initially suggesting target-site regeneration were attributed to *phes*\* mutations or truncations (Fig. S1G).

To test whether IS110 and bRNA alone are sufficient for transposition, we reconstituted transposition *in vitro* using purified *Caz*IS110-1 transposase, *in vitro*-transcribed bRNA, and plasmid substrates containing either donor/target sites (integration) or LT-RD/LD-RT junctions (excision) (Fig. 1I). Integration reactions yielded LT-RD and LD-RT joints exclusively with both components, as confirmed by PCR and Sanger sequencing. Excision reactions yielded rejoined donor-site (LD-RD) (Fig. 1J). These results demonstrate that *Caz*IS110-1 is an active transposon, and that *Caz*IS110-1 with its bRNA is sufficient to catalyze both excision and insertion without additional cofactors. Notably, excision rejoins the donor-site but leaves the target-site unjointed.

Since the primary evolutionary function of an active transposon is to proliferate within host genomes, the previously proposed cut-out-paste-in mechanism poses a risk of permanent loss unless a mechanism exists to preserve the original copy. Thus, we next investigated the transposition mechanism of *Caz*IS110-1 to understand how it ensures its proliferation.

### *Caz*IS110-1 excises without evidence of canonical cut-out-paste-in mechanism

To track the fate of the original transposon post-excision, we generated *E. coli* SX001 containing a linearized pRSF-1b plasmid flanked by LT-RD and LD-RT sites (referred to as the cargo). Excision of the cargo produced a functional, circular pRSF-1b plasmid, allowing kanamycin-resistant colony formation (Fig. 2A). After induction of *Caz*IS110-1 and its bRNA in SX001, we observed kanamycin-resistant colonies. Colony PCR and Sanger sequencing confirmed precise excision and formation of the LD-RD donor site (Fig. 2B and S2A). Mutation of the conserved E60 in the N-terminal DEDD motif or S312 in the C-terminal Tnp domain abolished excision, confirming essential roles of the RuvC-Tnp composite active site.

**Figure 2.**
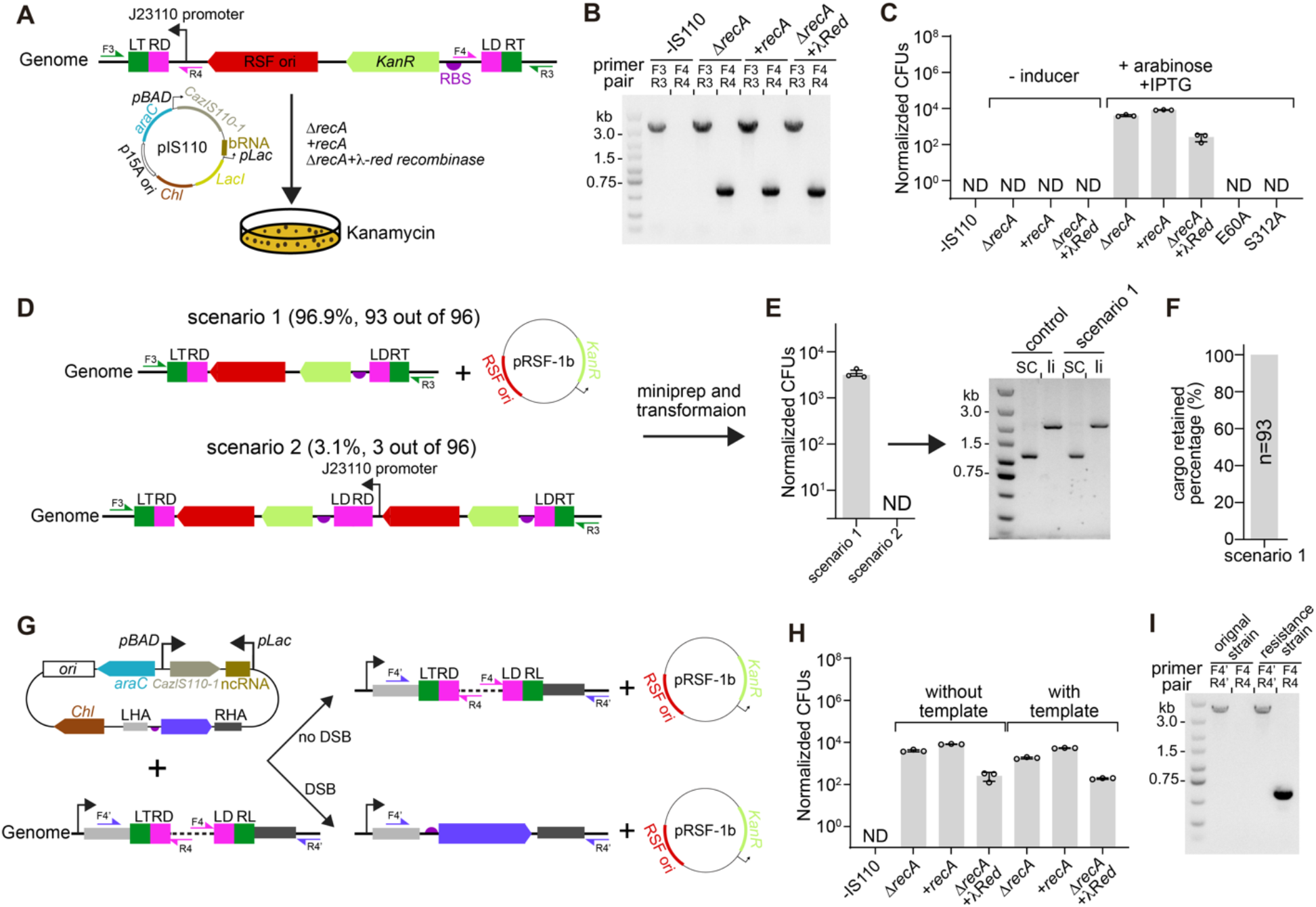
*Caz*IS110-1 does not use a cut-out-paste-in mechanism for its transposition. **A**, Experimental workflow to track the transposition *in vivo*. A linearized pRSF-1b plasmid flanked by LT-RD and LD-RT was inserted in *lacZ*. Excision yields a circular pRSF-1b plasmid conferring kanamycin resistance. **B**, PCR analysis of kanamycin-resistant colonies to assess donor site regeneration under the indicated conditions. **C**, Quantification of kanamycin-resistant colonies from **A** under indicated conditions. **D**, Kanamycin-resistant colonies belongs to two scenarios: scenario 1, excision yielded a circular pRSF-1b plasmid while retaining the original linearized cargo; scenario 2, a tandem cargo repeat. **E**, Plasmids from kanamycin-resistant colonies for scenario 1 and 2 were extracted and re-transformed into *E. coli* Top10. Only scenario 1 colonies yielded extractable plasmid, verified by plasmid linearization. SC: supercoiled; li: linearized. **F**, Quantification of the retained linearized pRSF-1b cargo at the original site in scenario 1 colonies (n=93). **G**, Experimental workflow for transposon fate with a homologous repair template. **H**, Quantification of kanamycin-resistant colonies from **G** under indicated conditions. **I**, PCR analysis of colonies from **G** for cargo retention and donor site regeneration. Data in **C, E**, and **H** are mean ± s.d. for n=3 biologically independent experiments.

Next, we investigated the fate of the original cargo in kanamycin-resistant colonies and observed two colony types (Fig. 2D). In ∼96.9% of cases (93/96), excision produced a circular pRSF-1b plasmid, with the original cargo retained in 100% of these cells (Fig. 2D-F). In ∼3.1% of cases (3/96), a tandem cargo repeat formed, conferring kanamycin resistance (Fig. 2D and S2B). This repeat suggested a possible mechanism for preserving the original cargo while simultaneously producing a functional circular copy. To determine whether tandem repeats actively contribute to cargo preservation, we continuously induced the *Caz*IS110-1 system in tandem repeat strains and plated cells on media with or without kanamycin. If tandem repeats were intermediates of transposition (Fig. S2C), they would be expected to resolve into a circular, kanamycin-resistant plasmid while restoring the original cargo (Fig. S2D). However, all tandem repeats reverted to the original cargo configuration without producing additional circular copies (Fig. S2E-F), indicating tandem repeats are transient byproducts, not functional intermediates. These results suggest that IS110 transposase actively preserves the original cargo during excision but not through tandem cargo repeat formation.

If IS110 utilized a canonical cut-out-paste-in mechanism, the non-rejoined target site left after excision would generate a DSB, potentially repaired by homologous recombination using the sister chromosome as a template. This process could restore the original cargo and serve as a potential preservation mechanism. To test this, we performed the excision assay in a Δ*recA* strain (Fig. 2A). Excision efficiency was reduced ∼2-fold in the absence of RecA, and complementation with the Lambda-Red recombinase system failed to restore excision efficiency (Fig. 2B-C), ruling out homologous recombination as the primary repair pathway.

We further tested this potential DSB-driven repair by providing an exogenous homologous repair template with 500-bp arms adjacent to the excision site (Fig. 2G). If homologous recombination were involved, this template should enhance cell survival. However, excision efficiency remained unchanged, and no insertion of the template into the cargo site was detected (Fig. 2H). All kanamycin-resistant strains retained the linearized pRSF-1b copy at its original genomic site (Fig. 2I). These findings suggest that *Caz*IS110-1 excision proceeds through a mechanism that avoids DSB formation, consistent with a non-cut-out mode of transposition.

### *Caz*IS110-1 employs a copy-out-paste-in mechanism

Our data do not support the use of the cut-out-paste-in mechanism by *Caz*IS110-1. The presence of a non-rejoined target-site after excision suggests that IS110 employs a replicative copy-out-paste-in mechanism, where only one DNA strand is transferred without rejoining the target-site. There are three DNA-based replicative transposition mechanisms^14-17^: cointegrate formation (e.g., Mu, Tn3), replication-dependent donor junction rejoining (e.g., *Helitrons*), and replication-independent donor junction rejoining (e.g., IS911) (Fig. S3). Cointegrate formation involves a Shapiro intermediate and does not regenerate the donor junction^18^, excluding this model. The second model requires replication machinery for donor junction rejoining^19^, whereas *Caz*IS110-1 and bRNA catalyze donor junction formation independently of additional cofactors (Fig. 1I), supporting the third, replication-independent mechanism.

To determine the excised strand, we performed an *in vitro* junction formation assay using oligonucleotides with phosphorothioate (PS) modifications at LT-RD and LD-RT^20^, on either the top or bottom strand (Fig. 3A). PCR showed that PS modifications reduced but did not abolish donor site rejoining (Fig. 3B). qPCR revealed that top-strand modification strongly inhibited rejoining, whereas bottom-strand modification had no negative effect (Fig. 3C). Dual-strand modification yielded the same inhibition as top-strand modification, indicating that the IS110 transposase excises and rejoins the top-strand. To confirm this, we developed a strand-specific PCR assay (Fig. 3D). Donor junction products were detected exclusively with top-strand-specific primers (Fig. 3E). Donor junction formation was strictly dependent on bRNA, as neither TBL-only nor DBL-only RNA supported the reaction. Together, these results demonstrate that IS110 employs a copy-out mechanism, excising and rejoining the top-strand.

**Figure 3.**
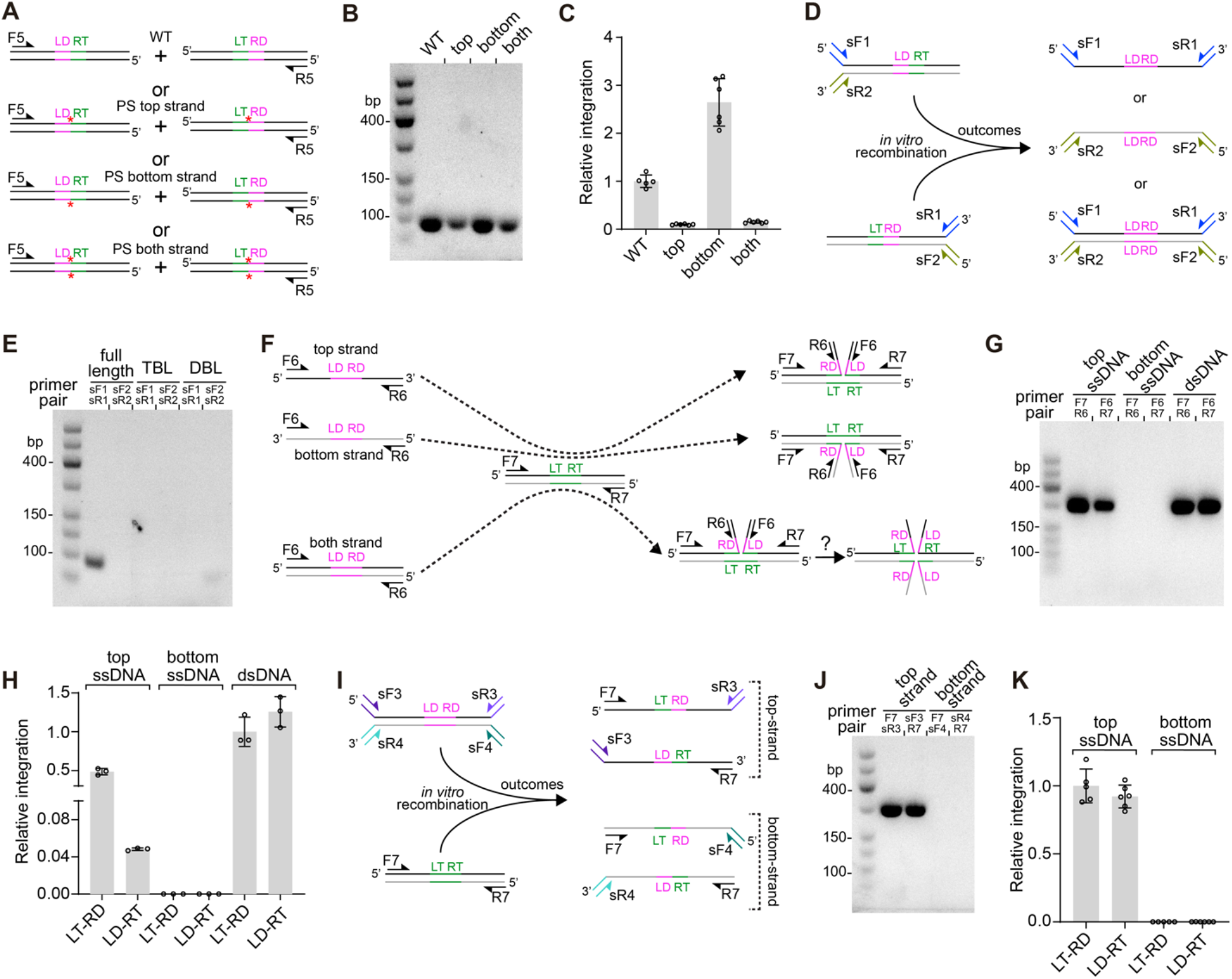
*Caz*IS110-1 catalyzes excision, donor rejoining, and integration via the top strand. **A**, Schematic of the excision and donor rejoining assay using phosphorothioate (PS)-modified substrates. Five central nucleotides at the LT-RD and LD-RT junctions were PS-modified on top-strand, bottom strand, or both. PS-modified strand was indicated with a red star. **B**, PCR analysis of reaction mixtures from **A. C**, qPCR-based quantification of donor site regeneration in reaction mixtures from **A**, comparing the effects of top- or bottom-strand PS modifications. **D**, Schematic of strand-specific PCR assay to determine rejoining strand. Primer pairs specific for top or bottom strand were indicated. **E**, PCR analysis of reaction mixtures from **D. F**, Schematic of donor/target integration assay using single-stranded (top or bottom) or double-strand donors. **G**, PCR analysis of reactions from **F. H**, qPCR-based quantification of integration efficiency across indicated donor types in **F. I**, Schematic of strand-specific PCR assay to assess strand preference during integration. Primer pairs specific for top or bottom strand were indicated. **J**, PCR analysis of reaction mixtures from **I. K**, qPCR-based quantification of nLT-RD and LD-nRT junctions formation on each strand for reaction mixtures from **I**. Data in **C, H**, and **K** are mean ± s.d. for n=3 biologically independent experiments.

The rejoined top-strand can be resolved into either a ssDNA or dsDNA circular intermediate. To test whether both forms integrate, we performed a PCR based integration assay (Fig. 3F). Both top-strand ssDNA and dsDNA integrated efficiently, while bottom-strand ssDNA did not (Fig. 3G). For the LT-RD site, dsDNA and top-strand ssDNA showed similar efficiency, but for the LD-RT site, dsDNA was ∼20-fold more efficient (Fig. 3H), indicating a substrate preference for dsDNA.

Hiraizumi et al. proposed that IS110 transposase-mediated the integration follows the canonical Holliday junction model^6^, involving top-strand cleavage/exchange, junction formation, and bottom-strand resolution. To test this, we applied strand-specific PCR to integration products (Fig. 3I). Surprisingly, integration occurred via top-strand insertion into the top-strand of the target, with equal efficiency in forming LT-RD and LD-RT junctions. Increasing PCR cycles revealed a faint bottom-strand LT-RD ligation product, but no LD-RT product (Fig. 3J). qPCR confirmed that top-strand integration was >1,300-fold more efficient than bottom-strand integration (Fig. 3K). These findings suggest that integration primarily occurs through the top-strand, indicating a potential alternative to the previously proposed Holliday junction-mediated dual-strand mechanism. Completion of full integration may require additional factors that facilitate bottom-strand joining.

### *Caz*IS110-1 employs an alternative RNA-guided “direct-transfer” mechanism

Our results demonstrate that both circular intermediate formation and integration require full-length bRNA, as neither TBL-only nor DBL-only RNA supports these processes. However, Siddiquee et al. reported TBL-only RNA-mediated transposition into preferred targets^7^, suggesting an alternative transposition pathway. We proposed that, instead of rejoining donor-site through top-strand excision and ligation at LD-RT and LT-RD, IS110 transposases can directly transfer top-strand from either junction to a new target (nLT-nRT), bypassing donor-site rejoining. This RNA-guided direct-transfer may be mediated by either bRNA or TBL-only RNA.

To test this, we employed strand-specific PCR and qPCR quantification (Fig. 4A). Only top-strand transfer products were detected, bottom-strand was not transferred in any condition. Both bRNA and TBL-only RNA supported LT-RD to nLT transfer, forming the nLT-RD junction, though bRNA was ∼5-fold more active (Fig. 4C). Given that TBL-only RNA is ∼10-fold more abundant *in vivo* (Fig. 1D), its overall contribution to this step is comparable to that of bRNA. In contrast, only bRNA catalyzed efficient LD-RT to nRT transfer, yielding the LD-nRT junction, while TBL-only RNA showed ∼1024-fold lower activity and DBL-only RNA was inactive in both reactions (Fig. 4B and C), indicating that LD-RT to nRT transfer is dependent on full-length bRNA. When comparing the overall efficiency of the two transfer steps, both bRNA and TBL-only RNA contribute to the LT-RD to nLT reaction, resulting a ∼10-fold more active than LD-RT to nRT step. This supports a sequential mechanism in which LT-RD to nLT transfer precedes LD-RT to nRT. Additionally, although LT-RD and LD-RT excision (copy-out pathway) mediated by bRNA was ∼1.8 fold more efficient than LT-RD to nLT transfer (direct-transfer) (Fig. 4C), the higher abundance of TBL-only RNA and its specific activity in the direct-transfer pathway suggest that the direct-transfer and copy-out-paste-in pathways contribute comparably to IS110 propagation.

**Figure 4.**
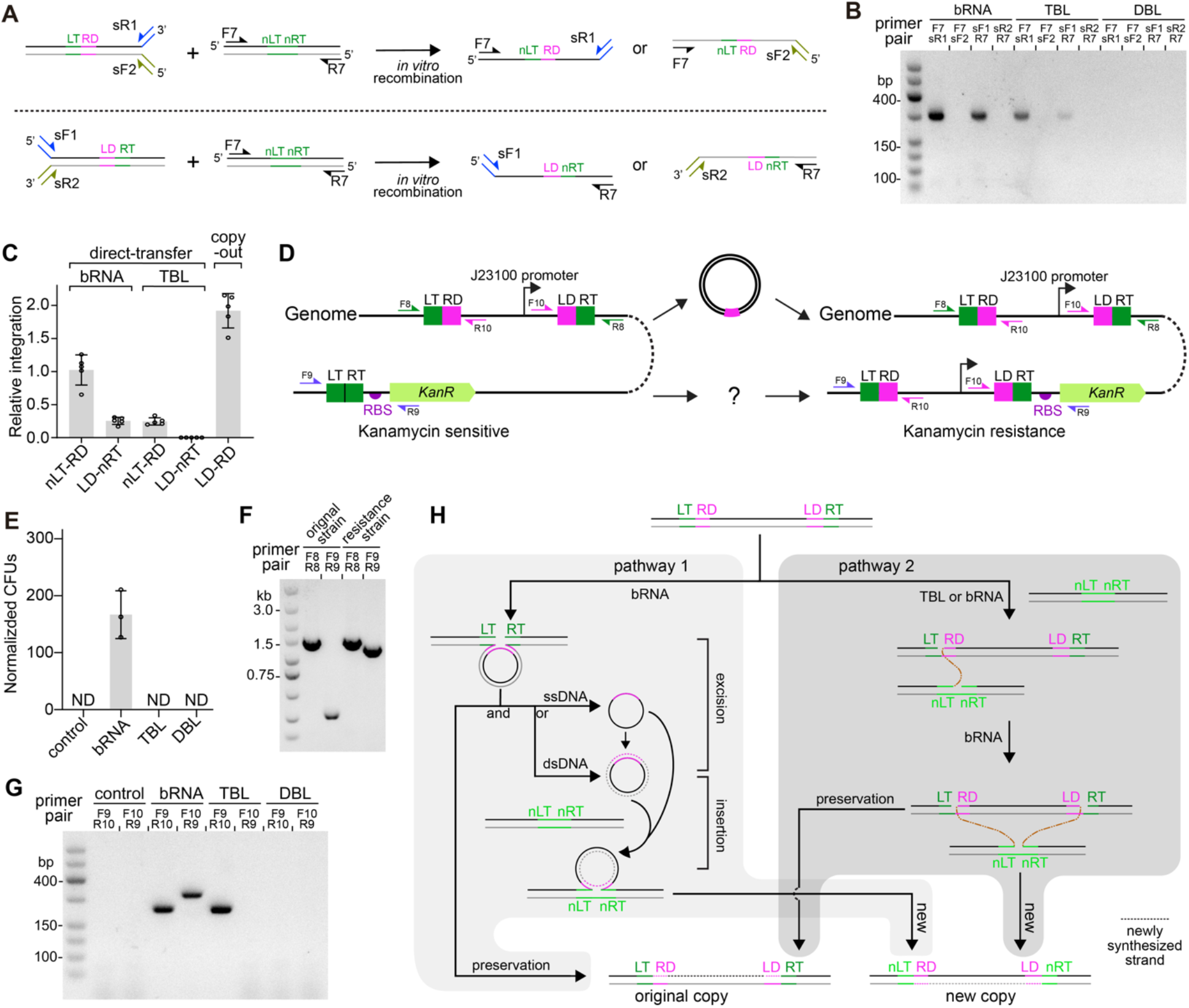
The direct-transfer mechanism of *Caz*IS110-1. **A**, Schematic of strand-specific PCR assays to detect recombinant junctions formed via direct transfer. Substrate design for detecting LT-RD junction (top) or LD-RT junction (bottom) transfer to a new target site, forming nLT-RD or LD-nRT. **B**, PCR analysis of reactions from **A. C**, qPCR-based quantification of direct transfer efficiency from **A. D**, Schematic of *in vivo* transposition assay. *E. coli* SX002 carries a mini-cargo with a J23110 promoter flanked by LT-RD and LD-RT, and a promoter-less *kanR* gene adjacent to the target. Successful transfer restores kanamycin resistance. **E**, Quantification of kanamycin-resistance colonies from **D** under indicated RNA conditions. **F**, PCR analysis of kanamycin-sensitive and -resistant colonies from **E. G**, PCR analysis of recombinant nLT-RD and LD-nRT junctions in cellular lysates. **H**, Model of dual RNA-guided transposition pathways. The first resembles a copy-out-paste-in mechanism using bRNA, where the IS110 transposase excises and rejoins the top-strand of the transposon, leaving the bottom strand intact. This is then resolved into single-stranded or double-stranded circular DNA. Both forms serve as substrates for top-strand integration into new target sites, while the bottom-strand remains unintegrated. In the second pathway, the transposase, with either TBL-only RNA or bRNA, catalyzes direct top-strand transfer from LT-RD to a new target site (nLT-nRT), forming the nLT-RD junction. Then, bRNA together with the transposase, but not TBL-only RNA, enables top-strand transfer of LD-RT to nRT, forming the LD-nRT junction. Both transposition modes create a new transposon copy while preserving the original. Data in **C** and **E** are mean ± s.d. for n=3 biologically independent experiments.

To test this process *in vivo*, we generated *E. coli* SX002 carrying a *lacZ*-interrupting mini-cargo flanked by LT-RD and LD-RT, along with a promoter-less kanamycin resistance gene downstream of *uidR* (Fig. 4D). Successful mini-cargo transposition into the target restores kanamycin resistance. We transformed SX002 with pIS110 plasmids expressing *Caz*IS110-1 and either bRNA, TBL-only RNA, or DBL-only RNA. Only bRNA-expressing cells formed kanamycin-resistant colonies. Colony PCR and Sanger sequencing confirmed correct mini-cargo transposition and demonstrated genome-wide transposition without strong locus preference (Fig. 4E-F and S4A-C). To directly assess intermediates formation, we performed PCR on cellular lysates. Both bRNA and TBL-only RNA supported formation of the nLT-RD junction, confirmed by Sanger sequencing (Fig. 4G and S4D), while the LD-nRT junction was detected only with bRNA. These results are consistent with the *in vitro* findings (Fig. 4B-C).

Although these data show that bRNA supports complete mini-cargo transposition *in vivo*, they do not distinguish the copy-out-paste-in and direct-transfer pathways, as bRNA mediates both. TBL-only RNA, which cannot support both excision and integration in the copy-out-paste-in pathway, should in principle isolate the direct-transfer pathway. However, due to its extreme inefficiently in catalyzing the second transfer step (LD-RT to nRT), no kanamycin-resistant colonies were recovered, leaving the direct transfer pathway unconfirmed *in vivo*. To address this, we turned to IS621, a IS110-family transposase characterized by Durrant et al.^5^, that shares the same RNA architecture. Using the same *in vivo* system, IS621 mediated efficient bRNA-driven transposition and, importantly, rare but detectable full transposition events with TBL-only RNA, confirmed by Sanger sequencing (Fig. S4E). Although >10,000-fold less efficient than bRNA, this provides direct *in vivo* evidence for the RNA-guided direct-transfer mechanism within the IS110 family.

Together, these findings reveal that *Caz*IS110-1 employs a distinct RNA-guided “direct-transfer” mechanism, in which the transposase sequentially transfers LT-RD and LD-RT directly to target sites, bypassing circular intermediate formation. TBL-only RNA supports partial transposition via LT-RD transfer, complete cargo mobilization requires full-length bRNA for the LD-RT transfer.

## Discussion

The recently identified IS110-bRNA system offers a promising platform for DSB-free genome editing by directly coordinating donor and target DNA recognition and integration. However, its transposition mechanism remained unclear. Here, we addressed several fundamental mechanistic questions using *Caz*IS110-1 and revealed two distinct transposition pathways, which appear to differ from earlier proposed models (Fig. 4H).

The first is a copy-out-paste-in mechanism, wherein the IS110 transposase excises the top strand while leaving the bottom strand intact, likely generating a figure-of-eight intermediate that resolves into ssDNA or dsDNA circular intermediate for integration. While this resembles IS911-like elements^16^, it is unique in its RNA guidance. Regarding the integration step, prior studies proposed a Cre-like, Holliday junction-mediated dual-strand insertion model^6^. Our data instead suggest that top-strand integration predominates, with >1,300-fold higher efficiency than bottom-strand integration. These findings raise the possibility that other additional factors may contribute to completing the integration of the bottom strand. The second pathway bypasses circular intermediates by directly transferring the top-strand sequentially from the LT-RD and LD-RT to new targets, forming nLT-RD and LD-nRT junctions. TBL-only RNA supports nLT-RD formation, while full-length bRNA supports both. This mechanism minimizes the loss of circular intermediates and ensures efficient transposon propagation. Although the copy-out-paste-in mechanism may entail a risk of incomplete transposition, it likely offers an evolutionary advantage by enabling horizontal gene transfer to new hosts.

Together, these findings help reconcile prior discrepancies. TBL-only RNA-mediated transposition events reported by Siddiquee et al. are consistent with partial transposition via the direct-transfer mechanism^7^, while bRNA-mediated events described by Durrant et al. align with the copy-out-paste-in pathway described here^5,6^. Notably, dual transposition strategies have been observed in other transposon systems. For example, Tn7 utilizes TnsABC-TnsD for homing and TnsABC-TnsE for mobility^21^, while CRISPR-associated transposases (CASTs) support both crRNA-guided and -independent homing integration^22,23^.

In summary, our findings suggest that IS110 transposon function through two distinct, RNA-guided strategies: a copy-out-paste-in mechanism and a direct top-strand transfer pathway. These insights not only advance our understanding of IS110 element biology but also provide a mechanistic framework that may inform future development of programmable, RNA-guided genome editing tools.

## Acknowledgments

This study was supported by National Key R&D Program of China (No.2023YFC3402301, C.X.), Strategic Priority Research Program of CAS (grant no. XDB0960300, C.X.), Haihe Laboratory (No.22HHSWSS00024, C.X.), CAMS Innovation Fund for Medical Sciences (CIFMS) (2021-I2M-1-041, S.R). SZU Top Ranking Project [86000000210 to H.L.].

## Author contributions

X.S., C.Y., Z.C., L.L., X.Y., and C.X. conceived and designed experiments. X.S., C.Y., Z.C., L.L., X.Y., performed experiments. X.S., C.Y., and C.X. analyzed the data, and X.S., S.R., H.L. and C.X. wrote the paper.

## Materials and Methods

### Plasmid construction

Primers and plasmids used in this study are described in Table S1. In brief, *CazIS110-1* gene was synthesized at Tsingke Biotech, along with the non-coding region. For recombinant *Caz*IS110-1 expression and purification, the *CazIS110-1* gene was cloned into pSV272, encoding an N-terminal His_6_-MBP (maltose binding protein) tag, the predicted bRNA was cloned into pET52b. For *in vivo* transposition assay, the *CazIS110-1* gene downstream of pBAD promoter and bRNA downstream of pLac promoter were cloned into pACYCDuet-1 to generate pIS110 plasmid. The *recA* or Lambda Red recombinase system was cloned into pIS110 to generate pIS110-*recA* and pIS110-λRed. To generate the pCargo plasmids, *phes\** gene under the control of J23100 promoter flanked with LT-RD/LD-RT junctions, or LD-RT/LT-RD junctions, or target/donor sites was cloned into pSC101. All these plasmids were cloned using a combination of Gibson assembly, BsaI mediated restriction digestion-ligation, and round-the-horn PCR. Plasmids were cloned, propagated in either DH5α or TOP10 cells (NEB), purified using Miniprep Kits (Aidlab), and verified by Sanger sequencing (Tsingke Biotech)

### *E. coli* strains construction

All *E. coli* strains used in *in vivo* transposition assays were derived from DH5α and constructed using scar-less λRed-mediated chromosomal editing with the pKD46-Cas9-λRed plasmid and the pCDF-sgRNA-SacB plasmid^24^. Donor DNA fragments with ∼500-bp homology arms used for recombineering were generated using overlay PCR. *E. coli* DH5α carrying the pKD46-Cas9-λRed plasmid was induced with 20 mM arabinose at OD600 = 0.2 for 2 h, then harvested and prepared as electrocompetent cells. Donor DNA and pCDF-sgRNA-SacB were co-electroporate into these cells. After 2 h recovery, cells plated on agar with ampicillin and chloramphenicol. Colonies were screened and validated by PCR the following day. pKD46-Cas9-λRed was cured at 42 °C, and pCDF-target-SacB was counter-selected on 5% sucrose plates. The primers involved in this process are all presented in Table S1.

### Preparation of dsDNA used for *in vitro* transposition assay

All DNA oligonucleotides were synthesized by GenScript Biotech. The DNA sequences used in this study are listed in Table S1. Oligos were dissolved in ddH_2_O to 100 μM. Two complementary oligos were mixed at a 1:1 ratio in annealing buffer (10 mM Tris, pH 7.5, 50 mM NaCl, 1 mM EDTA). Then the mixture was incubated at 95°C for 5 min and slowly cooled to room temperature by turning off the heat block. Annealed dsDNA substrates were stored at −80°C for until use.

### RNA structures

Predicted secondary RNA structures were generated using the Mfold web server^25^. All parameters were kept at their default settings. The predicted structures with the lowest free energy were visualized and used for downstream analysis.

### Small RNA sequencing

To extract RNA associated with *Caz*IS110-1 for RNA sequencing, 12 μL of purified *Caz*IS110 RNP complex (∼30 μM) was incubated with 1 μL of DNase I (20 U/ μL, NEB) in DNase I reaction buffer (10 mM Tris-HCl, pH 7.6, 2.5 mM MgCl_2_, 0.5 mM CaCl_2_) and incubated at 37 °C for 30 min. Next, 1 μL of Proteinase K was added, and the mixture was incubated at 50 °C for 10 min. After protein digestion, 100 μL of DEPC-treated water was added, followed by 100 μL of phenol:chloroform:isoamyl alchohol (25:24:1). The upper aqueous was collected and further extracted twice with 100 μL of chloroform. RNA in the final aqueous phase was purified using the DR04-Oligo Purification Kit (Aidlab) and eluted with 30 μL DEPC water.

The eluted RNA was treated with CIP and T4 PNK in T4 RNA ligase buffer with RNase inhibitor at 37 °C for 60 min, then purified again using the DR04-Oligo Purification Kit. For 3’-end labeling, poly(A) tails were added using *E. coli* Poly(A) polymerase. Polyadenylated RNA was reverse transcribed into cDNA using the Hifair® III 1^st^ strand cDNA synthesis kit with a specific primer (5′-CGTTCTCATGGCTCACGCAATTTTTTTTTTTTTTTTTTTT-3′). To degrade the RNA after reverse transcription, 1 μL of 5 M NaOH was added, and the sample was incubated at 93 °C for 3 min. After cooling to room temperature, the reaction was neutralized with HCl. The resulting cDNA was purified and concentrated using the DR06-Universal microDNA cleanup kit. For 5’-end adaptor ligations, a DNA adaptor was prepared by annealing two complementary oligos (5′-CCAGGGCTGTCGCCACAATGNNN-3′ and 5′-CATTGTGGCGACAGCCCTGG-3′). 1 μL of purified cDNA was mixed with 1 μL of 45 μM adapter and 1 μL T4-DNA ligase in T4 DNA ligase buffer and incubated at 25 °C for 5 h. Then ligation products were purified using the DR06-Universal microDNA cleanup kit. The samples were sequenced on the DNBSEQ-T7 platform (BGI) using a paired-end 150 bp protocol.

### Analysis of small RNA sequencing data

Raw paired-end FASTQ files were subjected to quality filtering using fastp^26^. Cleaned FASTQ files were subsequently merged using FLASH. The number of sequences containing only the target sequence, only the donor sequence, or both were quantified from the merged FASTQ file using Python. Merged FASTQ files were then aligned to predicted bRNA sequence using BWA and SAMtools. Small RNA-seq coverage across bRNA region was visualized with ggplot2.

### *In vitro* RNA transcription

The oligonucleotides used for DNA templates generation are listed in Table S1. RNAs were transcribed *in vitro* using linear DNA templates. Templates were generated by PCR using forward primers containing a T7 promoter sequence and reverse primers containing target-specific sequences. *In vitro* transcription was performed using the T7 RNA polymerase (NEB) in a 20 μL reaction containing 1 μg of purified DNA template, 7.5 mM each NTP, 40 U RNase inhibitor. Reactions were incubated at 37 °C for 6 h. Following transcription, 1 μL of DNase I (2 U/μL, NEB) was added and incubated at 37°C for 30 min to remove DNA template. Transcribed RNA was purified using the AidQuick Oligo Purification Kit (Aidlab, DR0401) and quantified using a NanoDrop spectrophotometer. Purified RNA aliquots were stored at −80 °C.

### Protein purification

*Caz*IS110-1 protein and *Caz*IS110-1–bRNA RNP complex were expressed in *E. coli* BL21 (DE3) cells. For *Caz*IS110-1 protein expression, the pSV272-*Caz*IS110-1 plasmid was transformed into BL21 (DE3). For RNA complex expression, a modified version of pSV272-*Caz*IS110-1 (lacking the MBP tag) was co-transformed with the pET52b-bRNA plasmid. Transformed cells were plated on LB agar containing appropriate antibiotics and incubated overnight at 37 °C. The following steps were the same for both protein and RNP expression. A single colony was picked and cultured overnight in 20 mL LB medium with appropriate antibiotics. The overnight culture was then transferred to 2 L LB medium supplemented with the same antibiotics and grown at 37 °C until reaching an OD_600_ of ∼0.5. Protein expression was induced by adding 0.2 mM IPTG, and cultures were incubated overnight at 18 °C. Cells were harvested by centrifugation and lysed by sonication in buffer A (20 mM Tris-HCl, pH 7.5, 500 mM NaCl, 10 mM imidazole, 10% glycerol, and 1 mM PMSF). After centrifugation, the supernatant was applied to HisPur Ni-NTA affinity resin. The resin was washed with buffer A containing 25 mM imidazole, and proteins were eluted with buffer A containing 250 mM imidazole. The eluted protein was concentrated to 1 mL and further purified on a Superdex 200 column in buffer A without imidazole. Peak fractions containing protein were pooled, concentrated, flash frozen in liquid nitrogen and stored at −80 ºC until further use.

### *In vitro* transposition assay

To assembly the *Caz*IS110-1-RNA complex, 12 μM *Caz*IS110-1 was incubated with 3 μM RNA (bRNA, TBL-only RNA, or DBL-only RNA) in reaction buffer (50 mM Potassium Acetate, 20 mM Tris-acetate, 10 mM Magnesium Acetate, 100 µg/ml Recombinant Albumin, pH 7.9). The mixture was incubated at 37°C for 30 min. For the transposition reaction, the pre-assembled *Caz*IS110-1-RNA complex was added to a final concentration of 300 nM along with 15 nM of the DNA substrates. The reaction was carried out at 50 ºC for 2 h and initiated by the addition of the *Caz*IS110-1-RNA complex. The reaction was terminated by adding EDTA to a final concentration of 50mM. The primers used to verify that this reaction occurs are described in Table S1.

### qPCR quantification assay

qPCR analysis for *in vitro* transposition reaction mixtures was performed using primers listed in Table S1. 20 µL reactions containing 10 µL of Power SYBR Green Mix, 0.4 µL of 10 mM primer pair, 1 µL of tenfold-diluted *in vitro* transposition reaction mixtures, and 8.2 µL H_2_O were prepared. Reactions were performed on an Applied Biosystems 7500 system using the following thermal cycling parameters: polymerase activation and DNA denaturation (95 °C for 5 minutes), 45 cycles of amplification (95 °C for 10 s, 60 °C for 45 s, and terminal melt-curve analysis (65-95 °C in 0.5 °C per 5 s increments).

### *In vivo* plasmid based transposition assay

pIS110 was co-transformed with pCargo plasmids containing either *phes*\* flanked LT-RD/LD-RT junctions (for excision) or target/donor sites (for integration) into *E. coli* DH5α. Transformants were plated on LB agar supplemented with chloramphenicol and apramycin, and incubated overnight at 37 °C. Single colonies were inoculated into 1 ml of LB broth with the same antibiotics and cultured at 37 °C for 12 h. Subsequently, 10 µl of this overnight culture was transferred into 1 ml of LB medium containing chloramphenicol, apramycin, 20 mM arabinose and 0.2 mmol IPTG, and incubated at 37 °C for 24 h. Cells were harvested by centrifugation and washed twice with sterile water. A portion of the resuspended cells was lysed by heating at 98°C for 10 min, and the resulting lysate was used as the template for PCR. ∼10^6^ cells from the remaining suspension were plated on YEG agar containing chloramphenicol, apramycin, and 16mM *p*-cl-Phe. Colonies from YEG plates were picked, resuspended in 10 µL of sterile water, and lysed at 98°C for 10 min for used as templates in colony PCR. The expected PCR products were confirmed by Sanger sequencing. The primers used for PCR are listed in Table S1.

### *In vivo* linearized pRSF-1b cargo excision assay

pIS110, pIS110-*recA*, pIS110-λRed plasmid were individually transformed into *E. coli* SX001 which harbors a linearized pRSF-1b plasmid flanked by LT-RD and LD-RT sites in the *LacZ* locus. Transformants were plated on LB agar supplemented with chloramphenicol and incubated overnight at 37 °C. Single colonies were inoculated into 1 mL of LB broth with the chloramphenicol, 20mM arabinose and 0.2mM IPTG, and incubated at 42 °C for 8 h. Then, the entire culture was centrifuged, and the cell pellet was resuspended and plated onto LB agar containing kanamycin. Plates were incubated overnight at 37 °C. Individual kanamycin-resistant colonies were picked, resuspended in 10 µL of sterile water, and lysed at 98°C for 10 min. The resulting lysate was used as template for colony PCR.

For plasmid extraction, single colonies belonging to scenario 1 and 2 were inoculated into 2 mL LB broth containing kanamycin and cultured overnight at 37 °C. Then, plasmids were extracted using the PL03-High Pure Plasmid Mini Kit. Extracted products were electrophoresed on 1% agarose gel, and DNA bands around 1500bp were excised and purified using the DR01-Gel Extraction Kit. The recovered product was re-transformed into *E. coli* Top10 and plated on LB agar containing kanamycin, followed by overnight incubation at 37 °C. A single clone was picked up and cultured in 2mL of LB broth containing kanamycin overnight at 37 °C. Plasmids were extracted and digested with AvrII (NEB), and digestion products were resolved on a 1% agarose gel. The expected PCR products were confirmed by Sanger sequencing. The primers used for PCR are listed in Table S1.

### *In vivo* genome-based transposition assay

pIS110-*recA* with either full-length bRNA, TBL-only RNA, or DBL-only RNA was transformed into *E. coli* SX002, which carrying a *lacZ*-interrupting mini-cargo flanked by LT-RD and LD-RT and a promoter-less kanamycin resistance gene downstream of *uidR*. Transformants were plated on LB agar supplemented with chloramphenicol and incubated overnight at 37 °C. Single colonies were inoculated into 1 mL of LB broth with the chloramphenicol, 20mM arabinose and 0.2mM IPTG, and incubated at 42 °C for 8 h. Then, the entire culture was centrifuged, and the cell pellet was resuspended and plated onto LB agar containing kanamycin and incubated overnight at 37 °C. Individual kanamycin-resistant colonies were picked, resuspended in 10 µL of sterile water, and lysed at 98°C for 10 min. The resulting lysate was used as template for colony PCR. The expected PCR products were confirmed by Sanger sequencing. The primers used for PCR are listed in Table S1.

### Statistics and reproducibility

qPCR and analytical PCRs resolved by agarose gel electrophoresis yielded similar results in three independent replicates. Sanger sequencing of PCR products was performed at least once. RNA sequencing was conducted once. All other data are expressed as the mean ± standard deviation (SD) from at least three independent experiments.

## References

1. Siguier, P., Gourbeyre, E., Varani, A., Ton-Hoang, B., and Chandler, M. (2015). Everyman’s Guide to Bacterial Insertion Sequences. Microbiol Spectr 3, MDNA3-0030-2014. 10.1128/microbiolspec.MDNA3-0030-2014.

2. Fedoroff, N.V. (2012). Presidential address. Transposable elements, epigenetics, and genome evolution. Science 338, 758–767. 10.1126/science.338.6108.758.

3. Knott, G.J., and Doudna, J.A. (2018). CRISPR-Cas guides the future of genetic engineering. Science 361, 866–869. 10.1126/science.aat5011.

4. Pacesa, M., Pelea, O., and Jinek, M. (2024). Past, present, and future of CRISPR genome editing technologies. Cell 187, 1076–1100. 10.1016/j.cell.2024.01.042.

5. Durrant, M.G., Perry, N.T., Pai, J.J., Jangid, A.R., Athukoralage, J.S., Hiraizumi, M., McSpedon, J.P., Pawluk, A., Nishimasu, H., Konermann, S., and Hsu, P.D. (2024). Bridge RNAs direct programmable recombination of target and donor DNA. Nature 630, 984–993. 10.1038/s41586-024-07552-4.

6. Hiraizumi, M., Perry, N.T., Durrant, M.G., Soma, T., Nagahata, N., Okazaki, S., Athukoralage, J.S., Isayama, Y., Pai, J.J., Pawluk, A., et al. (2024). Structural mechanism of bridge RNA-guided recombination. Nature 630, 994–1002. 10.1038/s41586-024-07570-2.

7. Siddiquee, R., Pong, C.H., Hall, R.M., and Ataide, S.F. (2024). A programmable seekRNA guides target selection by IS1111 and IS110 type insertion sequences. Nat Commun 15, 5235. 10.1038/s41467-024-49474-9.

8. Partridge, S.R., and Hall, R.M. (2003). The IS1111 family members IS4321 and IS5075 have subterminal inverted repeats and target the terminal inverted repeats of Tn21 family transposons. J Bacteriol 185, 6371–6384. 10.1128/JB.185.21.6371-6384.2003.

9. Higgins, B.P., Carpenter, C.D., and Karls, A.C. (2007). Chromosomal context directs high-frequency precise excision of IS492 in Pseudoalteromonas atlantica. Proc Natl Acad Sci U S A 104, 1901–1906. 10.1073/pnas.0608633104.

10. Liu, G., Lin, Q., Jin, S., and Gao, C. (2022). The CRISPR-Cas toolbox and gene editing technologies. Mol Cell 82, 333–347. 10.1016/j.molcel.2021.12.002.

11. Mahmood, M.A., and Mansoor, S. (2023). PASTE: The Way Forward for Large DNA Insertions. CRISPR J 6, 2–4. 10.1089/crispr.2023.0001.

12. Wery, N., Moricet, J.M., Cueff, V., Jean, J., Pignet, P., Lesongeur, F., Cambon-Bonavita, M.A., and Barbier, G. (2001). Caloranaerobacter azorensis gen. nov., sp. nov., an anaerobic thermophilic bacterium isolated from a deep-sea hydrothermal vent. Int J Syst Evol Microbiol 51, 1789–1796. 10.1099/00207713-51-5-1789.

13. Miyazaki, K. (2015). Molecular engineering of a PheS counterselection marker for improved operating efficiency in Escherichia coli. Biotechniques 58, 86–88. 10.2144/000114257.

14. Curcio, M.J., and Derbyshire, K.M. (2003). The outs and ins of transposition: from mu to kangaroo. Nat Rev Mol Cell Biol 4, 865–877. 10.1038/nrm1241.

15. Hickman, A.B., and Dyda, F. (2016). DNA Transposition at Work. Chem Rev 116, 12758–12784. 10.1021/acs.chemrev.6b00003.

16. Chandler, M., Fayet, O., Rousseau, P., Ton Hoang, B., and Duval-Valentin, G. (2015). Copy-out-Paste-in Transposition of IS911: A Major Transposition Pathway. Microbiol Spectr 3. 10.1128/microbiolspec.MDNA3-0031-2014.

17. Shkumatov, A.V., Aryanpour, N., Oger, C.A., Goossens, G., Hallet, B.F., and Efremov, R.G. (2022). Structural insight into Tn3 family transposition mechanism. Nat Commun 13, 6155. 10.1038/s41467-022-33871-z.

18. Shapiro, J.A. (1979). Molecular model for the transposition and replication of bacteriophage Mu and other transposable elements. Proc Natl Acad Sci U S A 76, 1933–1937. 10.1073/pnas.76.4.1933.

19. Grabundzija, I., Hickman, A.B., and Dyda, F. (2018). Helraiser intermediates provide insight into the mechanism of eukaryotic replicative transposition. Nat Commun 9, 1278. 10.1038/s41467-018-03688-w.

20. Kennedy, A.K., Haniford, D.B., and Mizuuchi, K. (2000). Single active site catalysis of the successive phosphoryl transfer steps by DNA transposases: insights from phosphorothioate stereoselectivity. Cell 101, 295–305. 10.1016/s0092-8674(00)80839-9.

21. Faure, G., Saito, M., Benler, S., Peng, I., Wolf, Y.I., Strecker, J., Altae-Tran, H., Neumann, E., Li, D., Makarova, K.S., et al. (2023). Modularity and diversity of target selectors in Tn7 transposons. Mol Cell 83, 2122–2136 e2110. 10.1016/j.molcel.2023.05.013.

22. Petassi, M.T., Hsieh, S.C., and Peters, J.E. (2020). Guide RNA Categorization Enables Target Site Choice in Tn7-CRISPR-Cas Transposons. Cell 183, 1757–1771 e1718. 10.1016/j.cell.2020.11.005.

23. Saito, M., Ladha, A., Strecker, J., Faure, G., Neumann, E., Altae-Tran, H., Macrae, R.K., and Zhang, F. (2021). Dual modes of CRISPR-associated transposon homing. Cell 184, 2441–2453 e2418. 10.1016/j.cell.2021.03.006.

24. Jiang, Y., Chen, B., Duan, C., Sun, B., Yang, J., and Yang, S. (2015). Multigene editing in the Escherichia coli genome via the CRISPR-Cas9 system. Appl Environ Microbiol 81, 2506–2514. 10.1128/AEM.04023-14.

25. Zuker, M. (2003). Mfold web server for nucleic acid folding and hybridization prediction. Nucleic Acids Res 31, 3406–3415. 10.1093/nar/gkg595.

26. Chen, S., Zhou, Y., Chen, Y., and Gu, J. (2018). fastp: an ultra-fast all-in-one FASTQ preprocessor. Bioinformatics 34, i884–i890. 10.1093/bioinformatics/bty560.

