## Supplemental material for "IS110 transposon utilizes two mechanistically distinct RNA-guided transposition pathways"

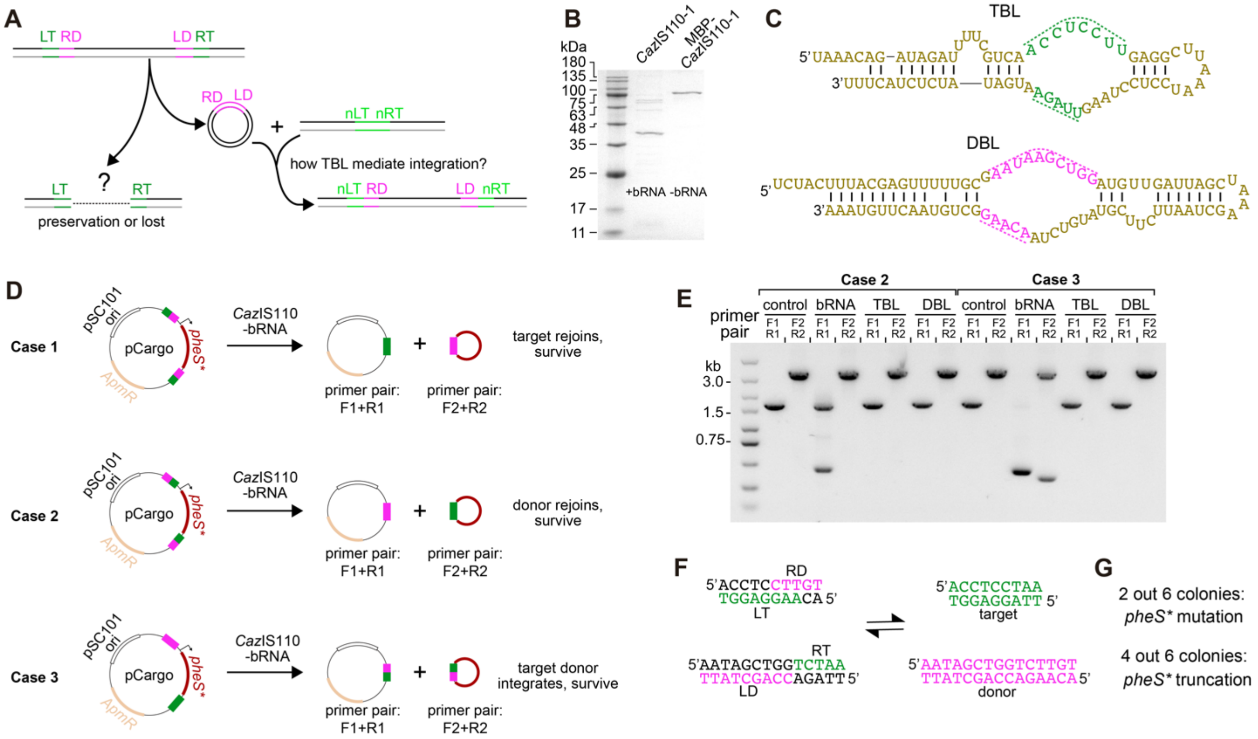


**Figure S1. *Caz*IS110 transposition features. Related to Figure 1. A,** Schematic of the transposition cycle of the IS110 elements. The elements are flanked with LT-RD and LD-RT junctions. IS110 elements are excised as circular dsDNA intermediates by rejoining LT-RD and LD-RT, restoring the donor-site (LD-RD), leaving the non-regenerated target sites. The role of the non-regenerated target sites in preserving the original IS110 element post-excision to ensure propagation remains unknown. Similarly, it is unclear whether and how TBL-only RNA, which lacks the donor DNA recognition loop, recognizes the donor DNA for integration into a target-site. **B,** SDS-PAGE of *Caz*IS110-1 with or without its predicted bRNA. **C,** Predicted secondary structure of the short RNA variants retaining only the conserved TBL (top) and DBL (bottom). **D,** Schematic of potential transposition outcomes in the three plasmid configurations used in **Fig. 1E**. Case 1, target site rejoining yields a functional *phes**-free plasmid; Case 2, donor site rejoining yields a functional *phes**-free plasmid; Case 3, integration between target and donor sites generates a functional *phes**-free plasmid. **E,** PCR analysis of transposition products from cells expressing *Caz*IS110-1 and either full-length bRNA, TBL-only, or DBL-only RNA. **F,** Schematic of excision and integration reactions, highlighting donor, target, LT-RD and LD-RT sequences. **G,** Colonies from Case 1, initially suggestive of target-site regeneration, were found to carry *phes** mutations or truncations, indicating that the resistance phenotype did not from true target-site rejoining.

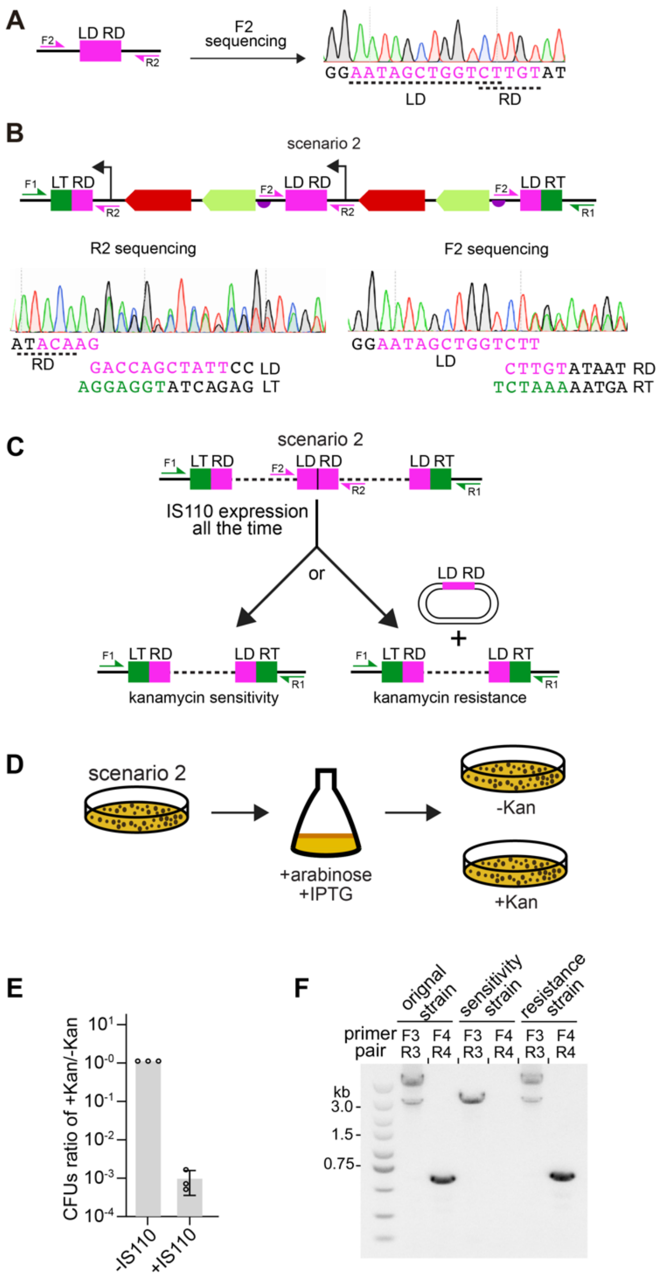


**Figure S2. Tandem cargo repeats are not functional intermediates in *Caz*IS110 transposition. Related to Figure 2. A,** Sanger sequencing of PCR products from a kanamycin-resistant strain confirms precise rejoining of the donor-site (LD-RD). **B,** Sanger sequencing of PCR products from a tandem cargo repeat strain reveals the formation of a precise donor site within the repeated structure. **C,** Schematic of hypothesized tandem repeat resolution outcomes. Route 1: the tandem repeat resolves back to its original single-copy configuration. Route 2: tandem repeat resolves into one circular pRSF-1b copy and one original single-copy configuration. **D,** Schematic of the tandem repeat resolution assay. A colony harboring a tandem repeat is induced to express *Caz*IS110-1 and full-length bRNA, followed by plating equal cell numbers on media with or without kanamycin. If resolution follows route 1, fewer colonies will form on kanamycin plates, as only unresolved tandem repeats confer kanamycin resistance. If resolution follows route 2, colony numbers will be similar on both plates. **E,** Quantification of colony formation on media with and without kanamycin from **D** under indicated experimental conditions. **F,** PCR and gel analysis of colonies from plates with or without kanamycin.

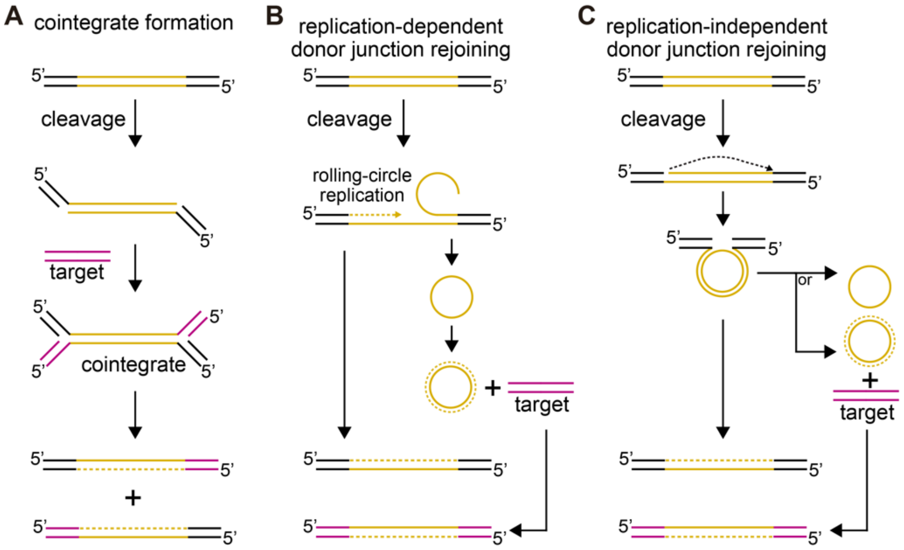


**Figure S3. Models of the copy-out-paste-in transposition mechanisms. Related to Figure 3. A,** Schematic of replicative transposition via cointegrate formation, which produces a Shapiro intermediate and does not rejoin the donor-site. **B,** Schematic of replication-dependent donor junction rejoining, where donor rejoining is coupled to host replication machinery. **C,** Schematic of replication-independent donor junction rejoining, where donor rejoins without requiring replication factors. Data from **Figure. 1L** support this third mechanism for *Caz*IS110-1, as excision occurs independently of additional cofactors and results in a rejoined donor junction.

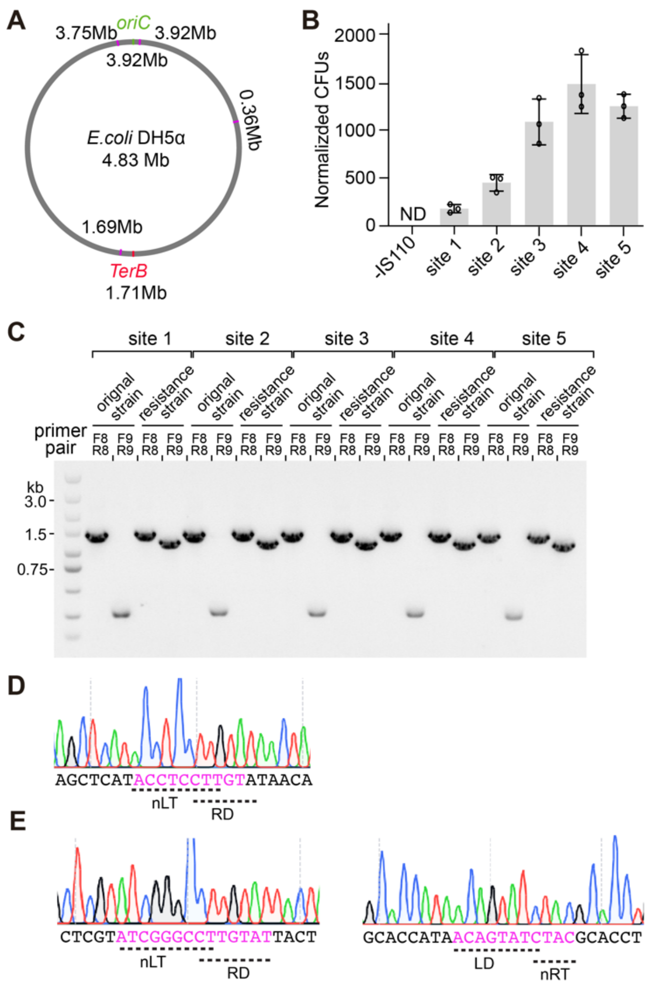


**Figure S4. Transposition efficiency of *Caz*IS110-1 at selected genome-wide target sites. Related to Figure 4. A,** Genome-wide distribution of the five selected target sites used to assess the *Caz*IS110-1 meditated transposition efficiency. **B,** Quantification of kanamycin-resistance colony formation at the five selected target sites with full-length bRNA. **C,** PCR and agarose gel analysis of kanamycin-sensitive and -resistant colonies from **B**. **D,** Sanger sequencing of the ligated nLT-RD PCR products from a lysate with TBL-only RNA of *Caz*IS110-1. **E,** Sanger sequencing of the ligated nLT-RD and LD-nRT PCR products from a kanamycin-resistant strain with TBL-only RNA of IS621.
